# Characterisation and manufacture of a *Neisseria gonorrhoeae* challenge agent for use in an oropharyngeal controlled human infection model

**DOI:** 10.64898/2026.08.06.743127

**Authors:** Georgina L Pollock, Shivani Pasricha, Kristy I. Azzopardi, Dana de Kretser, Evgeny A Semchenko, Kate L Seib, Joshua Osowicki, Deborah A Williamson, Eloise Williams, James S McCarthy

## Abstract

**Background:** Despite the importance of oropharyngeal gonorrhoea in transmission, suboptimal antimicrobial responses and propensity for horizontal transfer of antimicrobial resistance at this site, it remains understudied. An oropharyngeal *N. gonorrhoeae* controlled human infection model (CHIM) represents a promising tool to study infection and undertake translational research.

**Methods:** A panel of five contemporary *N. gonorrhoeae* isolates were subject to detailed characterisation to assess antimicrobial susceptibility, *in vitro* infectivity, cytotoxicity and serum sensitivity to inform challenge agent selection. A method for challenge agent manufacture, including release testing, was developed and validated.

**Findings:** All candidate isolates were able to infect the surface of pharyngeal and cervical cells *in vitro*. One isolate displayed an invasive phenotype, induced higher inflammatory cytokine production and displayed elevated serum resistance and was excluded. The remaining four isolates were minimally inflammatory, did not induce cytotoxicity and were susceptible to serum killing. Three of the four isolates grew in a defined liquid medium. Together these results led to the selection of a contemporary *N. gonorrhoeae* isolate suitable for use in CHIM. A challenge agent manufacture workflow was established and shown to reliably and reproducibly generate doses suitable for direct inoculation in an oropharyngeal CHIM.

**Conclusion:** Phenotypic characterization of candidate *N. gonorrhoeae* challenge agents led to the successful identification of a contemporary isolate suitable for implementation in a novel oropharyngeal gonorrhoea CHIM. We demonstrate the feasibility of a challenge inoculum manufacturing process that aligns with international best practice guidelines.

## INTRODUCTION

*Neisseria gonorrhoeae*, the causative agent of gonorrhoea, is a major global public health problem with an estimated incidence of 88 million cases per year [1, 2]. Infection typically occurs at mucosal sites including the genital tract, rectum and oropharynx. Complications include pelvic inflammatory disease, ectopic pregnancy and infertility, blinding newborn infection, and rarely, disseminated gonococcal infection [3].

Effective management of *N. gonorrhoeae* is threatened by the increasing prevalence of antimicrobial resistance (AMR), with emergence of resistance to all antimicrobials recommended for treatment [4]. Oropharyngeal gonorrhoea is also more difficult to treat, partly due to the unfavourable pharmacokinetics of many antibiotics in the oropharynx, resulting in a significant proportion of treatment failures at the oropharynx [5, 6]. While the new antimicrobials gepotidacin [7] and zoliflodacin [8] appear promising for the treatment of uncomplicated gonorrhoea, zoliflodacin is less efficacious for treatment of pharyngeal infection [8], and studies have been insufficiently powered to assess the efficacy of gepotidacin at extragenital sites [7, 9]. In addition, the oropharynx is increasingly being recognized as an important reservoir for *N. gonorrhoeae* transmission and acquisition of AMR [10–12].

Interest in developing an effective vaccine remains high, with several novel *N. gonorrhoeae*-specific vaccines currently under development [13]. However, in recently completed studies with vaccines against the closely related species *N. meningitidis,* no significant reduction in incidence of gonorrhoeae among men who have sex with men was reported [14, 15]. Together, these factors highlight the importance of investigating the efficacy of drugs, vaccines and other interventions at the oropharynx.

Controlled human infection models (CHIMs) provide a platform for investigating novel vaccines, therapeutics and other intervention strategies. Previous work established the feasibility and safety of the male urethral gonococcal CHIM using two laboratory adapted strains isolated in the 1970s, [16]. However, to date no oropharyngeal gonorrhoea CHIM has been developed. An oropharyngeal gonorrhoea CHIM could therefore assess the performance of novel drugs and interventions, as well as providing a model to study oropharyngeal infection [17].

We are developing an oropharyngeal gonorrhoea CHIM using a rational, transparent framework for the selection of a contemporary challenge agent, combined with a modernised approach to inoculum preparation and a carefully designed, community-informed protocol [18, 19]. In prior work, we described a systematic challenge-agent selection strategy that integrated available genomic and clinical metadata, guided by the dual principles of maximising participant safety and optimising model generalisability [20]. This process led to the shortlisting of five candidate *N. gonorrhoeae* isolates from an initial panel of 5,881 isolates. In the present study, we undertook detailed phenotypic characterisation of these five candidates to identify the strain most suitable for use. We also describe and evaluate the performance of a cell bank manufacturing approach including quality-control measures. Collectively, this work seeks to strengthen the safety, generalisability and reproducibility of an *N. gonorrhoeae* CHIM.

## METHODS

### Bacterial isolates and growth media

*N. gonorrhoeae* FA1090 was provided by Marcia Hobbs, University of North Carolina. Other *N. gonorrhoeae* isolates were provided by the Microbiological Diagnostic Unit Public Health Laboratory (MDU PHL, University of Melbourne). Desired colony types (Pili+ Opa-) were selected based on visual determination and confirmed by Western blot for Opa and scanning electron microscopy (SEM) for piliation. Bacterial isolates were cultured on gonococcal agar with Kellogg’s supplement (GCK) at 37°C, 5% CO_2_ overnight or in gonococcal base liquid with Kellogg’s (GCBL) or Graver-Wade liquid media (GW) where indicated.

### Mammalian cell culture

HeLa (CRM-CCL-2, ATCC) and Detroit 562 (CCL-138, ATCC) cells were maintained at 37°C, 5% CO_2_ in Minimum Essential Medium (MEM, Thermo Fisher Scientific) supplemented with non-essential amino acid solution (NEAA, 1X, Thermo Fisher Scientific), sodium pyruvate (1mM, Thermo Fisher Scientific) and 10% heat-inactivated foetal bovine serum (HI-FBS, Bovogen).Cells were seeded at ∼10^5^ cells/well in 24 well-plates the day prior, washed with PBS before bacterial suspensions (OD_600_ 0.005) were added (MOI ∼5-10). Infected cells were incubated for 1 hr at 37°C, 5% CO_2_. Immediately following infection, each inoculum was enumerated by 10-fold serial dilution in GCBL and spot plated onto GCK agar, before being incubated overnight at 37°C 5% CO_2_.To quantify attachment and invasion, cells were washed with PBS 1 hr after infection and media replenished before incubation for 5 hr. Media was then replaced with either fresh infection media (for ‘total attached’ quantification) or infection media containing gentamicin (20μg/mL) for 30 min (for ‘invaded’ quantification), then cells were washed again with PBS and lysed with GCBL-saponin (1% w/v) for 15 min. Lysates were collected and resuspended, before being serially diluted 10-fold in GCBL and spot-plated onto GCK agar for overnight incubation. Results were generated from at least three independent assays. ‘Total attached’ CFU were reported as mean CFU ± SEM as a percentage of the input inoculum, ‘invaded’ CFU are presented as mean CFU ± SEM as a percentage of ‘total attached’ CFU. Significance was calculated to compare each candidate isolate with FA1090 control using one-way ANOVA with Dunnett’s multiple comparison test.

### IL-8 and IL-6 cytokine ELISA

Cells were infected as described above. Following 1 hr infection, cells were washed with PBS to remove unattached bacteria, media was replaced with fresh media and incubated for a further 5 hr. Positive control wells were stimulated with TNF (20ng/mL, eBioscience). Following incubation, supernatant was collected and stored at -20°C prior to analysis using Human IL-8 and IL-6 ELISA kits (OptEIA, BD). Supernatants were generated from three independent experiments, each sample being tested in duplicate. Values are represented as mean ± SEM, significance was calculated using one-way ANOVA with Dunnett’s multiple comparisons test.

### Cytotoxicity

Cells were infected as described above. Following 1 hr attachment period, cells were washed with PBS to remove unattached bacteria, media was replaced with phenol red-free MEM (Thermo Fisher Scientific) supplemented with L-glutamine (2mM, Thermo Fisher Scientific), NEAA (1X, Thermo Fisher Scientific), sodium pyruvate (1mM, Thermo Fisher Scientific) and HI-FBS (5%, Bovogen) and incubated for a further 5 hr. Following incubation, supernatant was collected and assayed in triplicate using the CytoTox 96^®^ Non-radioactive cytotoxicity assay (Promega) as per manufacturer’s instructions. Results were generated from at least three independent assays. Absorbance values were blank subtracted and calculated as LDH activity as a percent of the positive control. Graphs show mean ± SEM, significance was calculated using one-way ANOVA with Dunnett’s multiple comparisons test.

### Scanning Electron Microscopy (SEM)

Isolates were grown overnight on GCK agar then suspended in GCBL at OD_600_ 0.1 and incubated on glass coverslips for 3 hr at 37°C. Cells were seeded onto glass coverslips then infected as described above. After 1 hr, cells were washed with PBS and media replenished. Cells were incubated for an additional 5 hr then fixed to glass coverslips with glutaraldehyde (2.5%, v/v in water) overnight.

Samples were washed three times with water before being dried by sequential 5 min submersions in ethanol of increasing concentration (50%, 70%, 80%, 90%, 95%, 100%, v/v in water) before dehydration using a critical point drier. Dried coverslips were mounted to stubs and sputter coated with gold (7 nm) then imaged using the FEI Teneo VolumeScope 3D SEM.

### Serum sensitivity assay

A pool of normal human serum (NHS) was generated by collecting blood samples from three healthy donors with no reported *N. gonorrhoeae* infection or meningococcal B vaccine within the previous three years (approved by Office of Research Ethics and Integrity, University of Melbourne, 2023- 20366-47468-8). Pooled serum was heat-inactivated at 56°C for 30 min (HI-NHS). Isolates were grown overnight on GCK agar at 37°C, 5% CO_2_, resuspended in pre-warmed RPMI 1640 medium. NHS and HI-NHS were 2-fold serially diluted (100% to 1.6%) in RPMI, then inoculated with ∼10^3^ CFU of each isolate (final serum concentration 90% to 1.4%), and incubated at 37°C, 5% CO_2_ for 1 hr. Then pre-warmed GCBL media was added, samples were spot plated in duplicate onto GCK agar, grown overnight and colonies enumerated. At least three independent experiments were undertaken; data are presented as a percentage survival versus no-serum control. For each serum dilution, unpaired t-tests were used to compare percent survival in NHS compared to HI-NHS. Results are presented as mean ±SEM.

### Growth curve

*Following* overnight grown on GCK agar, colonies was resuspended in hanks Balanced Salt Solution (HBSS), then inoculated into 10mL of GCBL or GW at OD_600_ 0.1, grown at 37°C with shaking at 180rpm before measurement at OD_600_. Density was compared using unpaired t-tests. Results are presented as mean ±SEM.

### Cell bank manufacture

Production of Working Cell Banks (WCB) and Dose Cell Banks (DCB) and stability testing is described in detail in Supplemental Material.

## RESULTS

### In vitro infection for assessment of attachment and invasion

*In vitro* attachment and invasion assays were performed to confirm that they retained infectivity during isolation, passage and storage, and to identify if any isolate displayed an invasive phenotype, relative to comparator strain FA1090. Two cell lines were used for the infection assays; HeLa cells, and Detroit 562 cells (derived from pharyngeal epithelium). Piliated, opacity-negative (Pili+Opa-) colonies were used for infections (Fig. S1a,b), with the exception of AUSMDU00015497 where Opa- colonies could not be isolated. In HeLa cells, all isolates attached with similar efficiency to FA1090, except for AUSMDU00053933, which infected cells with higher efficiency at both 1 hr and 6 hr post infection (Fig. 1a,b). The higher rate of infection for AUSMDU00053933 seen at 6 hr likely resulted from more efficient initial attachment rather than faster growth rate, as no differences were observed when comparing isolate average doubling time during infection (Fig. 1c). Gentamicin assay performed 6 hr infection showed that all isolates were minimally invasive, except AUSMDU00015497, where more intracellular bacteria were recovered post-infection than the FA1090 control (Fig. 1d). All isolates successfully formed microcolonies on the surface of the HeLa cells as observed by scanning electron microscopy (SEM) (Fig. 1e).

**Figure 1.**
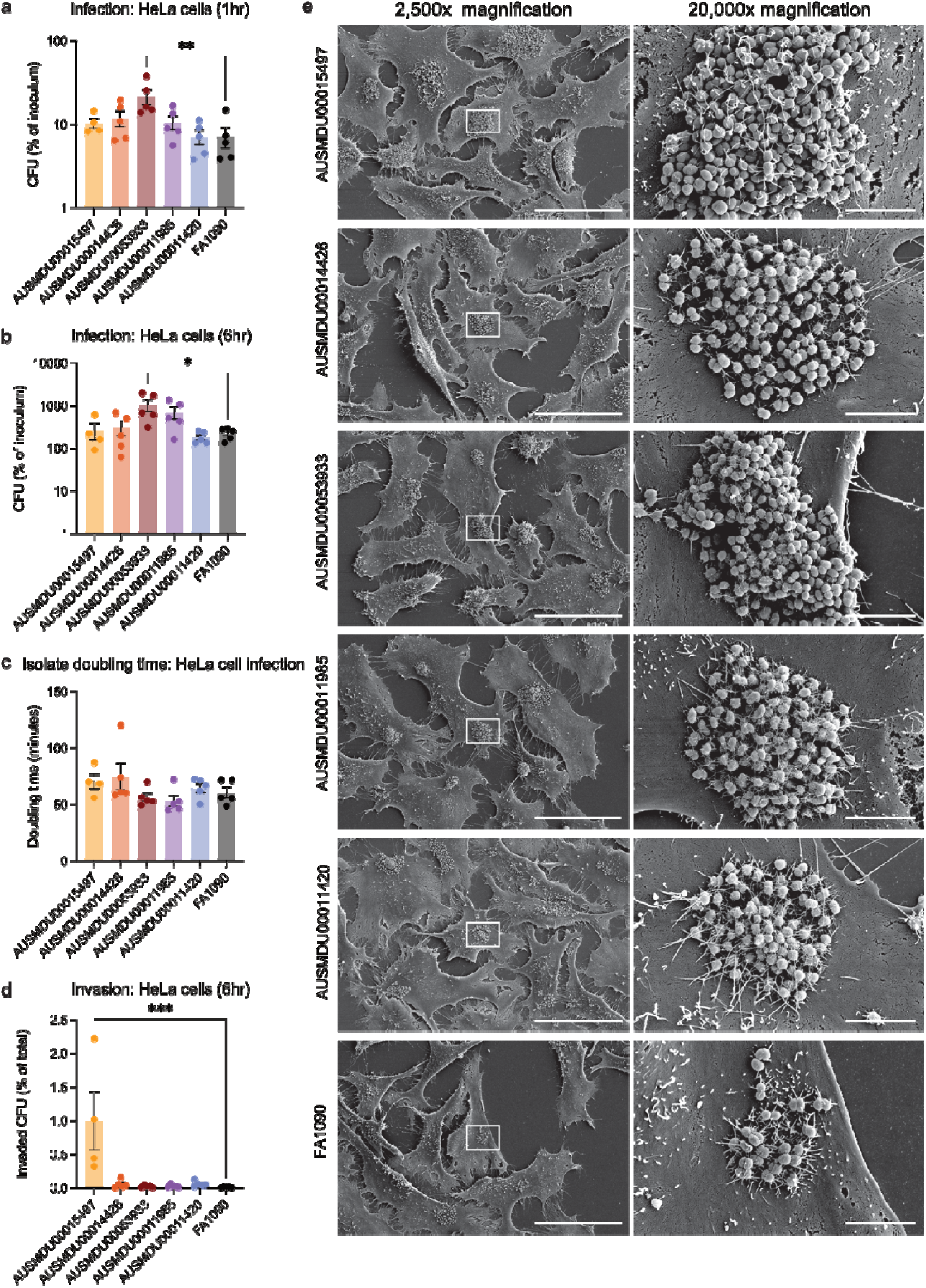
Candidate *N. gonorrhoeae* isolates infection of HeLa cells. Quantification of colony forming units (CFU) of candidate isolate strains following (a) 1hr or (b) 6hr infection of HeLa cells. Results are presented as a percentage of the input (inoculum). (c) Doubling time in minutes of each isolate during HeLa infection. (d) Quantification of intracellular CFU following 6hr infection of HeLa cells. Results are presented as a percentage of total CFU count at 6hrs. Filled circles represent individual datapoints from each independent experiment, bar and error represent mean ±SEM. Significance was calculated using one-way ANOVA with Dunnett’s multiple comparisons test, *P<0.05, **P<0.01, ***P<0.001, ns not significant. (e) Representative scanning electron micrographs of *N. gonorrhoeae* microcolonies following 6hr infection of HeLa cells imaged at 2,500x (scale bar 50μm) or 20,000x magnification (scale bar represents 5μm).

To our knowledge, Detroit 562 cells have not previously been used to model infection with *N. gonorrhoeae*. Similarly to HeLa cells, an increase in bacterial CFU from 1 hr to 6 hr post infection was observed, indicating that Detroit 562 cells can support *N. gonorrhoeae* replication (Fig. S2a,b). The infection efficiency and rate of growth on the host cell surface was consistent with that seen in HeLa cells. (Fig. S2c-e). Results of infection dynamics in Detroit 562 cells showed no significant difference in infectivity between the candidate isolates and FA1090 at 1 hr and 6 hr post-infection (Fig. 2a,b), nor were there strain specific differences in doubling time or invasion compared to FA1090 at 6 hr post infection (Fig. 2c,d). Microcolonies were also observed using SEM following 6 hr infection of Detroit 562 cells (Fig. 2e).

**Figure 2.**
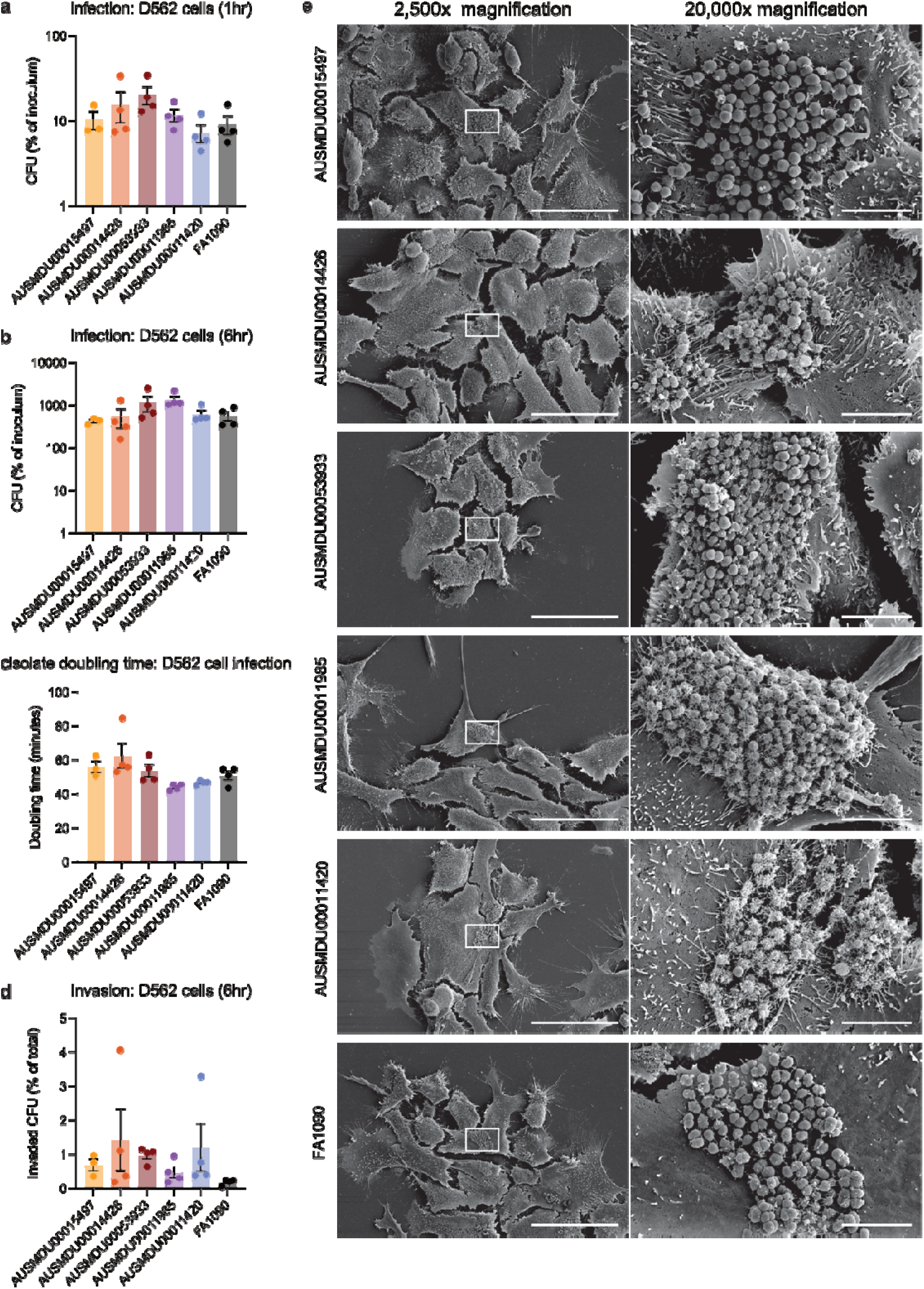
Candidate *N. gonorrhoeae* isolates infection of Detroit 562 cells. Quantification of colony forming units (CFU) of candidate isolate strains following (a) 1hr or (b) 6hr infection of Detroit 562 cells. Results are presented as a percentage of the input (inoculum). (c) Doubling time in minutes of each isolate during Detroit 562 cell infection. (d) Quantification of intracellular CFU following 6hr infection of Detroit 562 cells. Results are presented as a percentage of total CFU count at 6hrs. Filled circles represent individual datapoints from each independent experiment, bar and error represent mean ±SEM. Significance was calculated using one-way ANOVA with Dunnett’s multiple comparisons test. (e) Representative scanning electron micrographs of *N. gonorrhoeae* microcolonies following 6hr infection of Detroit 562 cells imaged at 2,500x (scale bar 50μm) or 20,000x magnification (scale bar represents 5μm).

### Host cell response to infection

Isolates were assessed for cytotoxicity of HeLa and Detroit 562 cell lines by lactate dehydrogenase (LDH) release assay compared to FA1090. Following a 6-hr incubation, LDH was measured in cell culture media as a measure of host cell death. Candidate isolates did not induce a cytotoxic response in HeLa or Detroit 562 cells compared to FA1090-infected or uninfected controls (Fig. 3a,b).

**Figure 3.**
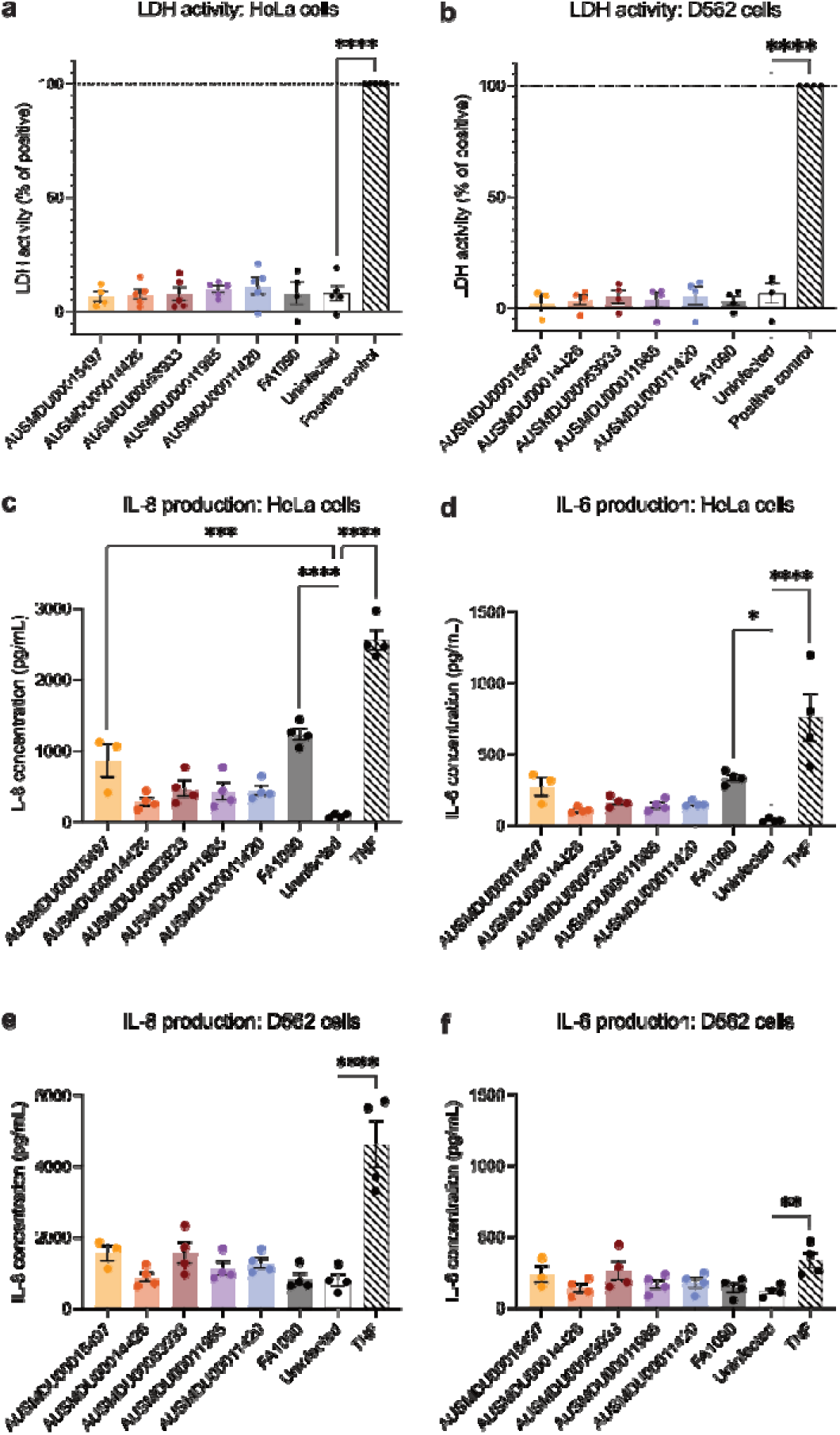
Host cell response following infection with candidate *N. gonorrhoeae* isolates. LDH activity in the culture supernatant of (a) HeLa or (b) Detroit 562 cells following 6hr infection with *N. gonorrhoeae* candidate isolates. Results are presented as LDH activity as percentage of the positive control (chemically lysed cells). Filled circles represent individual blank-subtracted datapoints from each independent experiment, bar and error represent mean ±SEM. Significance was calculated using one-way ANOVA with Dunnett’s multiple comparisons test, ****P<0.0001. Quantification of IL-8 and IL-6 production from HeLa cells (c-d) or Detroit 562 (e-f) following 6hr infection with candidate *N. gonorrhoeae* isolates. TNF stimulation was used as a positive control. Filled circles represent individual datapoints from each independent experiment, bar and error represent mean ±SEM. Significance was calculated using one-way ANOVA with Dunnett’s multiple comparisons test, *P<0.05, **P<0.01, ***P<0.001, ****P<0.0001.

Candidate isolates were assessed for their ability to induce the inflammatory cytokines IL-8 and IL-6 in HeLa and Detroit 562 cells as these are elevated in the urine and mucosal surfaces of men with gonorrhoea [21–23]. Infection with both AUSMDU00015497 and FA1090 led to an increase in IL-8 production compared to the other four isolates or uninfected cells (Fig. 3c). FA1090 alone induced IL- 6 production. In Detroit 562 cells no significant differences in IL-8 or IL-6 production were observed (Fig. 3e,f).

### Serum sensitivity

During our genomic-based strain selection [20], *porB1a*-possessing *N. gonorrhoeae* isolates were excluded due to association with DGI [24], likely mediated by resistance to killing conferred by binding to C4BP and factor H [25, 26]. To complement this genotypic assessment, survival was compared when incubated in either normal human serum (NHS) or heat-inactivated NHS (HI-NHS) across serum dilutions. *N. gonorrhoeae* WHO F was included as a PorB1a serum-resistant control isolate. As expected, *N. gonorrhoeae* WHO F showed equivalent survival in NHS and HI-NHS across all dilutions (Fig. 4a). FA1090 also displayed partial serum resistance consistent with previous reports (Fig. 4b)[25], with survival declining in serum concentration of ≥6.25%. only AUSMDU00015497, showed increased survival in up to 12.5% serum (Fig. 4c-g).

**Figure 4.**
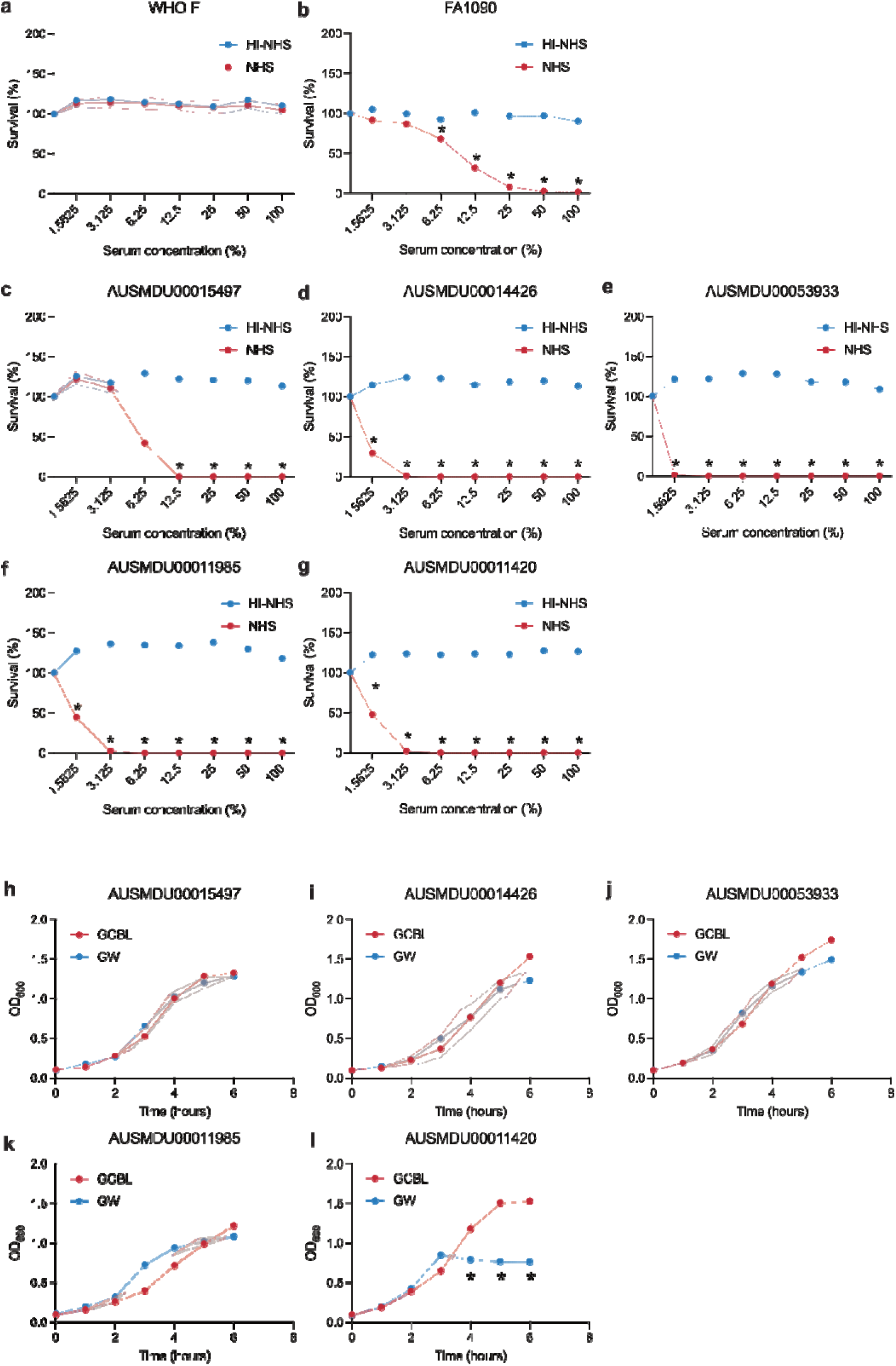
Serum sensitivity and growth of *N. gonorrhoeae* isolates. (a-g) Survival of *N. gonorrhoeae* candidate and control isolates in normal human serum (NHS, red) or heat-inactivated NHS (HI-NHS, blue). Filled circles represent mean survival expressed as a percentage of survival in no serum control. Shading represents SEM. Multiple unpaired *t*-tests were used to compare survival in NHS relative to HI-NHS control for each serum dilution. *P<0.05. (h-l) Growth curves of *N. gonorrhoeae* isolates in standard culture media GCBL (red) and chemically defined culture media GW (blue). Filled circles represent mean density (OD_600_), shading represents SEM. Multiple unpaired *t*-tests were used to compare density in GW relative to GCBL for each time point, *P<0.05.

### Growth in chemically defined medium

To maximise safety of the challenge inoculum for administration to human volunteers, we identified a chemically-defined growth media free from animal-derived products to reduce the risk of Transmissible Spongiform Encephalopathies (TSEs) [27]. We tested the suitability for supporting candidate isolate growth. Graver-Wade (GW) liquid medium [28] supported the growth of all candidate isolates except AUSMDU00011420 which failed to reach the same density and entered lag phase prematurely at 4 hr post-inoculation in GW (Figure 4h-l).

### Phenotypic antimicrobial susceptibility testing

Antimicrobial susceptibility testing was performed for all isolates to penicillin, spectinomycin, ceftriaxone, ciprofloxacin, tetracycline and azithromycin using the EUCAST method and MIC breakpoint interpretation (SupTable1). All isolates were classified as susceptible (or less susceptible in the case of penicillin) except AUSMDU00015497 which was resistant to tetracycline.

### Cell bank manufacture method development and validation

We adapted a pre-existing challenge agent cell bank manufacture strategy [29] to allow for extemporaneous administration of the challenge agent at bedside using single-use inoculum vials that could undergo rigorous release testing prior to use. (Figure 5a). Given the outcomes of phenotypic characterisation, two candidate isolates, AUSMDU00053933 and AUSMDU00011985, were tested in the cell bank manufacture process. Due to poor growth in the dose cell bank (DCB) phase of production (Fig. S3), AUSMDU00011985 was excluded, and AUSMDU00053933 was selected as the challenge agent (Table 1).

**Figure 5.**
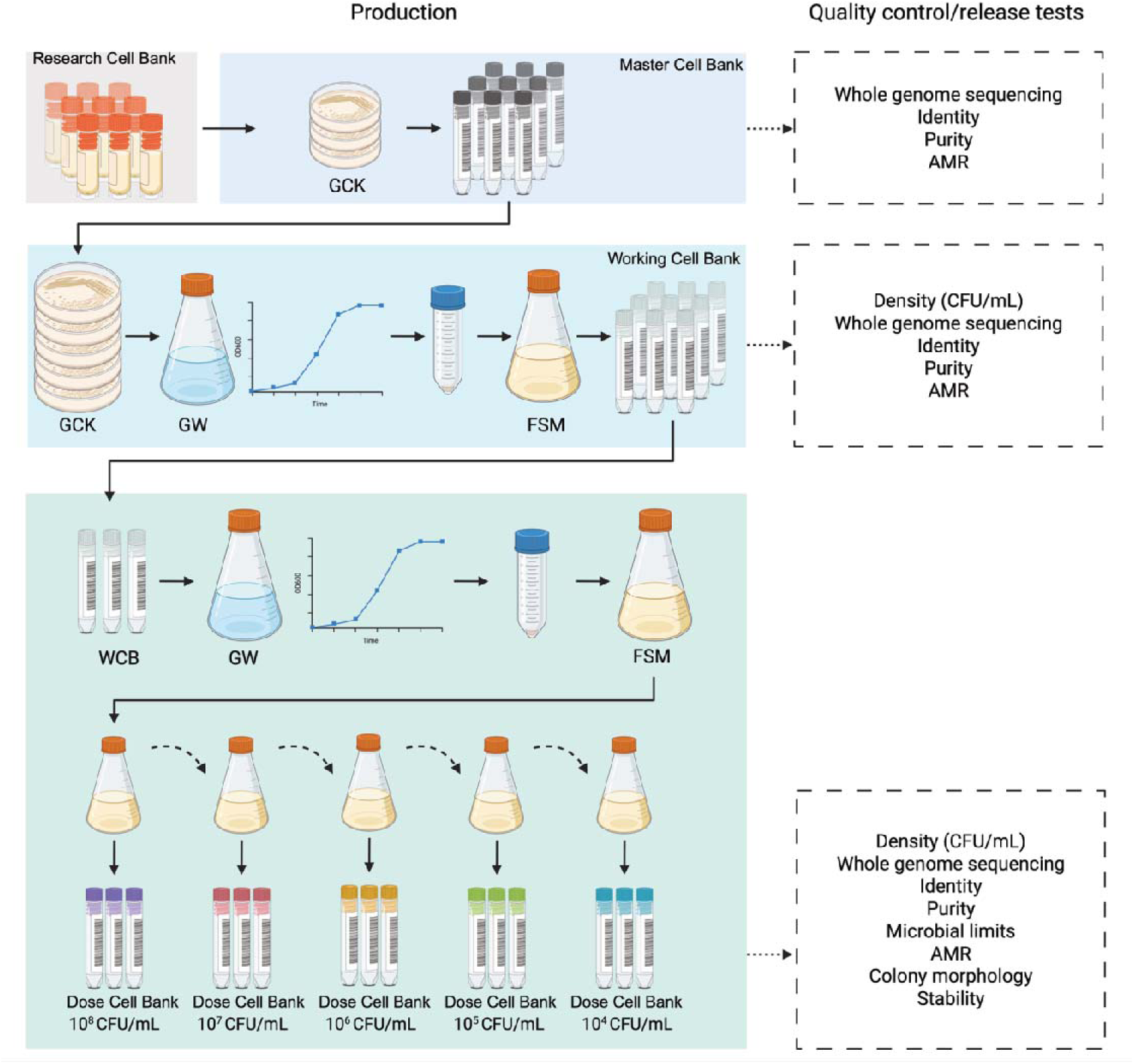
Cell bank production method and quality control. Schematic of the *N. gonorrhoeae* challenge agent cell bank production process and description of the quality control and release tests performed at each stage. GCK, gonococcal agar with Kelloggs supplements; GW, Graver Wade; FSM, Freezer Storage media; WCB, working cell bank; CFU/mL, colony forming units per millilitre. Figure created using a licensed copy of Biorender.

**Table 1.** Summary of candidate isolate phenotypic characterisation.

| Isolate | Year isolated | Anatomical site | Clinical notes | Lineage | MLST | Genomic AMR<sup>1</sup> | Phenotypic AMR | Attachment | Invasion | Cytotoxicity | Cytokine production | Serum Sensitivity | Growth in liquid | Manufacture |
| --- | --- | --- | --- | --- | --- | --- | --- | --- | --- | --- | --- | --- | --- | --- |
| AUSMDU00015497 | 2017 | Rectal | Asymptomatic (screening) | B | 1584 | Pass | TetR | Yes | Increased | No | Increased IL-8 | Decreased sensitivity | Yes | Not tested |
| AUSMDU00014426 | 2017 | Urogenital | Symptomatic | B | 1596 | Wild type | Pass | Yes | Minimal | No | No | Sensitive | Yes | Not tested |
| AUSMDU00053933 | 2020 | Urogenital | Symptomatic | B | 1596 | Wild type | Pass | Increased | Minimal | No | No | Sensitive | Yes | Pass |
| AUSMDU00011985 | 2017 | Oropharyngeal | Asymptomatic (screening) | B | 8122 | Pass | Pass | Yes | Minimal | No | No | Sensitive | Yes | Poor |
| AUSMDU00011420 | 2017 | Urogenital | Symptomatic | A | 1597 | Pass | Pass | Yes | Minimal | No | No | Sensitive | Poor | Not tested |
| FA1090 | 1970's | Endocervix | Probable DGI | B | 1899 | Pass | Pass | Yes | Minimal | No | Increased IL-8 & IL-6 | Decreased sensitivity | Not tested | Not tested |

DGI disseminated gonococcal infection. MLST multi locus sequence type. AMR antimicrobial resistance. <sup>1</sup>Genomic AMR as defined in [20]

Validation of manufacture was performed for isolate AUSMDU00053933 to generate three working cell banks (WCB1, WCB2 and WCB3), each of which was used to generate three Dose Cell Banks (nine in total; DCB1.1, 1.2, 1.3, DCB2.1, 2.2, 2.3 and DCB3.1, 3.2, 3.3). Key performance metrics included growth kinetics, pre- and post-thaw CFU, accuracy and reproducibility of doses generated, thaw stability and long-term freezer stability.

While variability in the rate of growth during the WCB liquid growth phase was observed between batches, all met the acceptable viability criteria of >10^7^ CFU/mL upon thawing (Fig. 6a-d). Similarly, variation was seen in the DCB liquid growth phase (Fig. 6e). Acceptance criteria for dose determination was defined as a post-thaw CFU of target dose ± 0.5log_10_. All dose levels across all generated DCBs met the criteria except DCB3.3 (dose 10^6^ CFU), demonstrating accuracy and reproducibility of the method (Fig. 6f-h). To ensure piliation was maintained colony phenotype was assessed by quantifying the number of piliated (small, domed, reflective) and non-piliated (large, flat) colonies. For all DCBs, piliation was >90% (Fig S4a). In an ongoing program to test long term stability of frozen dose vials, up to 10 months stability has been demonstrated (Fig 6q). Specifications for manufactured cell banks include: identification by MALDI-ToF, Gram stain, superoxol, oxidase and nucleic acid amplification, whole genome sequencing and typing, antimicrobial susceptibility testing, specific pathogen contamination testing, CFU enumeration and colony morphology, (Sup Table S2-4).

**Figure 6.**
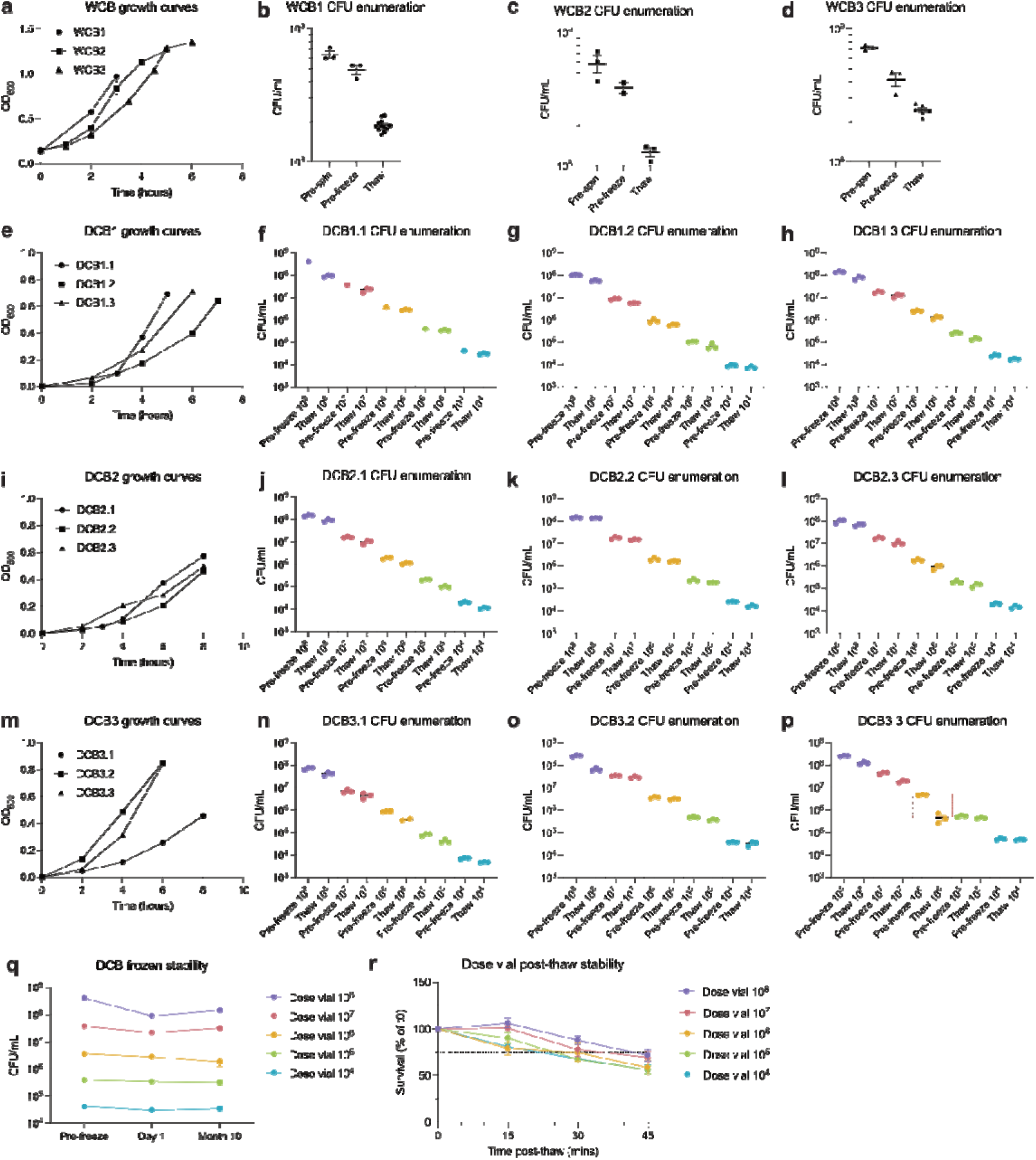
Cell bank manufacture validation. (a) Growth curves (OD_600_) of AUSMDU00053933 in the liquid growth phase of Working Cell Bank (WCB) manufacture for three WCB batches (WCB1, WCB2 and WCB3). (b-d) Colony forming units (CFU/mL) enumerated at the pre-spin, pre-freeze and thaw steps of WCB manufacture for three independent WCB batches (WCB1, WCB2 and WCB3), bar and error represent mean ±SEM. (e, i, m) Growth curves (OD_600_) of AUSMDU00053933 in the liquid growth phase of Dose Cell Bank (DCB) manufacture for nine independent DCB batches, grouped to correspond to the WCB from which they were derived. CFU enumeration for Dose Cell Bank batches DCB1.1-1.3 (f-h) DCB2.1-2.3 (j-l) and DCB3.1-3.3 (n-p) at the pre-freeze and thaw steps of manufacture for each target dose level, bar and error represent mean ±SEM. Boxed area indicates where acceptance criteria were not met. (q) Frozen dose stability (CFU/mL) over time for DCB vials, bar and error represent mean ±SEM. (r) Stability of thawed dose vials over time, bar and error represent mean ±SEM percentage survival compared to 0 minutes post-thaw.

### Inoculum stability and use

Dose stability upon thawing was assessed to inform dosing on the day of challenge. Single use inoculum vials were assessed for viability post-thaw for up to 45 minutes (Fig. 6r). Loss of viability was seen over time, informing the clinical practise of dose delivery within 15 minutes of thaw. To select a swab type for inoculum delivery, uptake and release of inoculum from three swab types (spun Rayon, flocked Nylon and flocked Polyester) was assessed. The flocked Nylon swab was selected for its higher and more consistent uptake compared to the flocked polyester swab (151mg±4.4mg versus 123mg±10.7mg, mean ±SD, respectively), and higher inoculum release than the spun rayon swab across all dose levels. The dose released from the swab was shown to be proportional to the dose vial inoculum, such that dose delivery increased proportionately with increasing dose vials (Fig. S4b,c).

## DISCUSSION

CHIMs are increasingly important tool for investigating the host pathogen interaction and for product development [30]. In this study, we describe the process undertaken to select and characterise a contemporary *N. gonorrhoeae* challenge agent for use in an oropharyngeal CHIM, and describe the process for its manufacture. To this end, we conducted phenotypic assessment of five contemporary *N. gonorrhoeae* candidate isolates, previously selected from a large pool using a genomics-based systematic approach based on the principles of maximising generalisability and safety [20]. This was based on criteria important for a challenge isolate, including retention of key infectivity determinants maximising study safety, and amenability to contemporary manufacture processes.

It is obvious that the selected challenge agent must retain infectivity with repeated passage and cryopreservation. Key phenotypic characteristics such as favourable colony types [31], attachment to host cells and formation of microcolonies are likely good surrogate markers, and involve membrane structures including pili, Opa, LOS, PorB, and NHBA [32, 33]. While the relative importance of each adhesin in the process of establishing human pharyngeal infection is unknown, our data indicate that the selected strain retains key colony phenotypes (Pili+ Opa-), preserved *in vitro* cell culture infection of both pharyngeal and cervical cells, and ability to form microcolonies.

Minimization of the risk of severe or invasive gonococcal infection such as DGI is essential. While determinants of invasive gonococcal infection are incompletely understood, they likely depend on both host and bacterial factors. The most well-defined bacterial correlate of invasive disease is the *porB1a* allele [24], likely due to the conferring serum resistance through binding to C4BP and Factor H [26, 34], and the ability to readily traverse epithelial cells [35]. A large majority (∼90%) of PorB1a isoforms can bind to C4BP, conferring serum resistance, but importantly, a small subset of PorB1b isoforms can do the same [36]. To minimise risk of severe disease in study participants, we had previously selected by bioinformatic analysis only *porB1b-*possessing candidate isolates [20]. Here we confirmed serum sensitivity, except AUSMDU00015497, which had a similar level of resistance to FA1090. Despite FA1090 being safely used to initiate experimental infection in hundreds of individuals [16], it was originally isolated from an individual with suspected DGI, and possesses a PorB isoform that confers serum resistance [37]. Host risk factors for invasive disease such as complement deficiency, can be controlled for by careful exclusion criteria in the study protocol [18].

In line with the guidance for challenge agent selection [27], we also characterised pathogenicity, including cytotoxicity and invasiveness and proinflammatory effect. All tested isolates were minimally invasive, except AUSMDU00015497. Notably, we were unable to isolate an Opa- variant of AUSMDU00015497. The increased propensity for invasion in HeLa cells of this isolate may potentially be due to expression of Opa, which has been reported to aid invasion of epithelial cells [38]. AUSMDU00015497 also induced IL-8 production following infection of HeLa cells, though no more so than FA1090. No isolate showed cytotoxicity. While these assays contribute to the characterisation of the *N. gonorrhoeae* isolate *in vitro*, we acknowledge the limitations of using cultured cells to predict clinical outcomes in the vastly more complex environment at human mucosal surfaces. Other facets of pathogenicity controlled by phase variation [39] were not modelled here.

In addition to characterising a novel challenge isolate, we demonstrated the feasibility of producing challenge inoculum bank aligned with the *Smart Practices for Production of Challenge Agents* guidance [40]. Thus we implemented measures to optimise quality control during manufacture. The current practise for inoculum preparation for *N. gonorrhoeae* CHIM entails overnight culture on solid media, followed by resuspension to the desired density (as determined by OD), and administration [41]. The need for immediate use of the inoculum using this approach limits quality control. Our approach, using a scalable, tiered cell bank structure results in production of single-use, cryopreserved vials at pre-determined doses. These require no further preparation at the study site before use. This approach also allows implementation of a quality control program that employs in-process and release testing at all stages of manufacture to confirm identity, purity and potency. In addition to improved safety, this approach improves reproducibility across multiple cohorts, studies, study arms and clinical sites.

In summary, we have presented a systematic and transparent approach to the selection and manufacture of a contemporary *N. gonorrhoeae* challenge agent. The infectivity, safety and tolerability of the *N. gonorrhoeae* AUSMDU00053933 manufactured challenge inoculum will be tested in a clinical trial [18] with the goal of establishing a robust and reliable model of oropharyngeal gonorrhoea to aid in the development of novel therapeutics and vaccines, and further our understanding of this complex organism.

## ACKNOWLEDGMENTS

Authors wish to acknowledge the facility staff, in particular Zlatan Trifunovic, at the Ian Homes Imaging Centre, Bio21 Molecular Science and Biotechnology Institute, University of Melbourne for their contribution to the electron microscopy in this study. We wish to thank the staff at the Microbiological Diagnostics Unit Public Health Laboratory for performing antimicrobial susceptibility testing for this study and Jim Ackland for Regulatory advice. All authors report no conflict of interest. The genome sequence of the isolate selected for manufacture can be retrieved from the ncbi archive (www.ncbi.nlm.nih.gov/biosample/?term=AUSMDU00053933).

## FUNDING AND CONTRIBUTIONS

This research was funded by the Medical Research Future Fund Clinician Researchers Applied Research in Health Grant (MRFAR000354). EW was supported by an NHMRC Postgraduate scholarship (2005380); KLS is supported by a NHMRC Investigator Grant (2017383). JO is supported by an NHMRC fellowship (2009548). JSM is supported by an NHMRC Investigator Grant (2016396).

## Notes

### Competing Interest Statement

The authors have declared no competing interest.

