## supplementals for "Characterisation and manufacture of a *Neisseria gonorrhoeae* challenge agent for use in an oropharyngeal controlled human infection model"

**SUPPLEMENTARY MATERIALS**

**SUPPLEMENTARY METHODS**

*Method for Cell Bank Manufacture*

To prepare the mock Working Cell Banks (WCBs), ten GCK agar plates were inoculated with N. gonorrhoeae isolate AUSMDU00053933 and grown for 16-20 hr at in a humidified incubator at 37°C, 5% CO_2_. Colonies were collected and resuspended in Graver Wade medium (GW). This suspension was used to inoculate 200 mL of GW at OD600 = 0.15 in a conical flask. The flask was incubated at 37°C, 200rpm, and sampled periodically to monitor density, until mid-log phase was reached. A sample of the culture was removed for enumeration of viable bacteria (“pre-spin” enumeration), with the remainder pelleted by centrifugation. Bacterial pellets were resuspended in freezer storage medium (FSM), pooled in a new conical flask, and volume restored to the original volume collected in FSM. A subsample of this suspension was removed for enumeration of viable bacteria (“pre-freeze” enumeration). The suspension was aliquoted into 1mL pre-labelled cryovials. These were submerged into a dry-ice alcohol bath to snap-freeze then transported to an ultra-low temperature freezer for storage.

To prepare mock Dose Cell Banks (DCB), vials of the WCB were thawed rapidly at 37°C, pooled and used to inoculate GW to a total volume of 120mL, in a conical flask. The flask was incubated at 37°C, 200rpm, and sampled periodically to monitor optical density, until the required density was reached. The culture was then centrifuged to pellet bacteria, and the bacterial pellets thoroughly resuspended in FSM. Doses were generated from this suspension by performing 10-fold serial dilution in FSM. A subsample of each suspension was removed for enumeration of viable bacteria (“pre-freeze” enumeration), then each dose suspension was aliquoted in 0.5 mL pre-labelled cryovials that were frozen as above.

To enumerate the WCB and DCB bank post-thaw, vials were rapidly thawed at 37°C and enumerated by serially diluting cultures in pre-warmed hanks Balanced Salt Solution (HBSS), then plated onto GCK agar in duplicate and incubating overnight at 37°C, 5% CO2 for colony enumeration the following day.

*Post-thaw dose stability assay*

To assess dose stability post-thaw, triplicate DCB vials were rapidly thawed and incubated at 37°C. At 0, 15, 30 and 45 min post thaw, dose vials were sampled and serially diluted in pre-warmed HBSS, then spot-plated onto GCK agar in duplicate. Cultures were incubated overnight at 37°C, 5% CO2 for enumeration. Results are presented as mean +/-SEM of triplicate vials, enumerated in duplicate, for each nominal dose level.

*Antimicrobial susceptibility testing*

Antimicrobial susceptibility testing was performed by the Microbiological Diagnostic Unit Public Health Laboratory (MDU PHL) using the EUCAST method and MIC breakpoint interpretation.

*Immunoblotting and PAGE*

To detect Opa expression, *N. gonorrhoeae* isolates were grown overnight on GCK agar then suspended in gonococcal base liquid with Kellogg’s (GCBL) at OD_600_ 0.1. Suspensions were pelleted by centrifugation and resuspended in water, before being heated at 95°C for 20 minutes. Lithium dodecyl sulfate (LDS) sample buffer (NuPAGE, Thermo Fisher Scientific) supplemented with dithiothreitol (DTT, 50mM) was added to samples before heating again at 70°C for 10 minutes, and then samples were resolved on 4-12% Bis-Tris Gels (Thermo Fisher Scientific) by PAGE. Proteins were transferred to nitrocellulose membranes which were blocked in skim milk solution (5% in PBS-Tween 0.05%) before being probed overnight with either anti-opa 4B12 or anti-H.8 (2-8C-4-1) (1:200 in skim milk solution) followed by anti-mouse-HRP (1:3000 in skim milk). Immunoblots were developed using Clarity ECL Western Blotting Substrates (BioRad). The 4B12 and 2-8C-4-1 monoclonal antibodies were obtained from the Developmental Studies Hybridoma Bank (DSHB), created by the NICHD of the NIH and maintained at The University of Iowa, Department of Biology, Iowa City, IA 52242.

*Swab uptake and release*

Three swab types were assessed for inoculum release, Spun Rayon (8155CIS, Copan), Flocked Nylon (552C, Copan) and Flocked Polyester (520CS01, Copan). Swabs were immersed in thawed dose vials for 10 seconds, before being immersed in 500μL HBSS for 10 seconds with gentle swirling. HBSS suspensions were mixed by pipetting, then diluted and spot plated onto GCK overnight for enumeration. To assess swab uptake amount and consistency, mock dose vials were thawed, then weighed before and after a swab submersion. Swab uptake amount was determined by change in vial weight following swab submersion.

**SUPPLEMENTARY TABLES**

**Table S1. Antimicrobial susceptibility testing results for candidate isolates**

| Isolate | Penicillin | | Spectinomycin | | Ceftriaxone | | Ciprofloxacin | | Tetracycline | | Azithromycin | |
| --- | --- | --- | --- | --- | --- | --- | --- | --- | --- | --- | --- | --- |
|  | MIC | Interpretation | MIC | Interpretation | MIC | Interpretation | MIC | Interpretation | MIC | Interpretation | MIC | Interpretation |
| AUSMDU00015497 | 0.5 | Less susceptible | ≤32 | Susceptible | ≤0.008 | Susceptible | ≤0.03 | Susceptible | 1 | Resistant | 0.125 | Likely susceptible |
| AUSMDU00014426 | 0.03 | Susceptible | ≤64 | Susceptible | ≤0.008 | Susceptible | ≤0.03 | Susceptible | ≤0.25 | Susceptible | 0.125 | Likely susceptible |
| AUSMDU00053933 | 0.06 | Less susceptible | ≤32 | Susceptible | ≤0.008 | Susceptible | ≤0.03 | Susceptible | ≤0.25 | Susceptible | ≤0.06 | Likely susceptible |
| AUSMDU00011985 | 0.125 | Less susceptible | ≤32 | Susceptible | ≤0.008 | Susceptible | ≤0.03 | Susceptible | 0.5 | Susceptible | ≤0.06 | Likely susceptible |
| AUSMDU00011420 | 0.25 | Less susceptible | ≤32 | Susceptible | ≤0.008 | Susceptible | ≤0.03 | Susceptible | 0.5 | Susceptible | ≤0.06 | Likely susceptible |

MIC: Minimum inhibitory concentration

**Table S2. Master Cell Bank product specification sheet**

| **Test category** | **Test** | **Accreditation** | **Sample tested** | **Amount of sample tested (minimum)** | **Specifications** |
| --- | --- | --- | --- | --- | --- |
| In process test (identity/purity) | Colony morphology | Not accredited | Research cell bank culture | All plates inspected | Circular, smooth, grey to white colonies. |
|  | Purity (visual) |  | Research cell bank culture | All plates inspected | No visible contamination |
| Release tests: Culture identification | MALDI | ISO 15189 | Master Cell Bank | 3 colonies/MCB vial | *N. gonorrhoeae* |
|  | Gram stain |  | Master Cell Bank | 3 colonies/MCB vial | Gram negative diplococci with no visible contamination |
|  | Superoxol test |  | Master Cell Bank | 3 colonies/MCB vial | Positive |
|  | Oxidase test |  | Master Cell Bank | 3 colonies/MCB vial | Positive |
| Release tests: AST characterisation | Antimicrobial susceptibility:  Penicillin  Spectinomycin  Ceftriaxone  Ciprofloxacin  Tetracycline  Azithromycin | ISO 15189 | Master Cell Bank | 3 colonies/MCB vial | Penicillin: susceptible or less susceptible  Spectinomycin: susceptible  Ceftriaxone: susceptible  Ciprofloxacin: susceptible  Tetracycline: susceptible  Azithromycin: likely susceptible |
| Release tests: Genomic identification and characterisation | Whole genome sequencing | Not accredited | Master Cell Bank gDNA | gDNA extract from MCB vial | MLST: 1596  NG-STAR: 4332  NG-MAST: (*tbpB* 3072, *porB* 7697) |
| Release test: material sterility | Total aerobic microbial count (TAMC) | Accredited for Appendix XVI B, British Pharmacopeia. | FSM | 1mL/lot | <1CFU/mL |
|  | Total Yeast and Mold count (TYMC) |  | FSM | 1mL/lot | <1CFU/mL |
|  | Microbial contamination test |  | FSM | 1mL/lot | Absence of bile-tolerant Gram negatives, Staphylococcus aureus, Pseudomonas aeruginosa, Escherichia coli, and Salmonella |

MCB: Master Cell Bank. AST: Antimicrobial Susceptibility Test. gDNA: Genomic DNA. FSM: Freezer storage medium. CFU: Colony forming unit. ISO: International Organisation for Standardisation.

**Table S3. Working Cell Bank product specification sheet**

| **Test category** | **Test** | **Accreditation** | **Sample tested** | **Amount of sample tested (minimum)** | **Specifications** |
| --- | --- | --- | --- | --- | --- |
| In process test (identity/purity) | Colony morphology | Not accredited | Master cell bank culture | All plates inspected | Circular, smooth, grey to white colonies. |
|  | Purity (visual) |  | Master cell bank culture | All plates inspected | No visible contamination |
| Release test: Density | CFU enumeration | Not accredited | Final culture with cryopreservative | Enumerated in technical triplicate | Report only |
|  |  |  | Final container Working Cell Bank | 3 vials (beginning, middle, end of vial fill) | ³ 10^7^ CFU/mL |
| Release tests: Culture identification | MALDI | ISO 15189 | Final container Working Cell Bank | 3 colonies/WCB vial | *N. gonorrhoeae* |
|  | Gram stain |  | Final container Working Cell Bank | 3 colonies/WCB vial | Gram negative diplococci with no visible contamination |
|  | Superoxol test |  | Final container Working Cell Bank | 3 colonies/WCB vial | Positive |
|  | Oxidase test |  | Final container Working Cell Bank | 3 colonies/WCB vial | Positive |
| Release tests: AST characterisation | Antimicrobial susceptibility:  Penicillin  Spectinomycin  Ceftriaxone  Ciprofloxacin  Tetracycline  Azithromycin | ISO 15189 | Final container Working Cell Bank | 3 colonies/WCB vial | Penicillin: susceptible or less susceptible  Spectinomycin: susceptible  Ceftriaxone: susceptible  Ciprofloxacin: susceptible  Tetracycline: susceptible  Azithromycin: likely susceptible |
| Release tests: Genomic identification and characterisation | Whole genome sequencing | Not accredited | Working Cell Bank gDNA | gDNA extract from WCB vial | MLST: 1596  NG-STAR: 4332  NG-MAST: (*tbpB* 3072, *porB* 7697) |
| Release tests: Microbiological limits and material sterility | Total aerobic microbial count (TAMC) | Accredited for Appendix XVI B, British Pharmacopeia. | Final container Working Cell Bank | 1mL | Report only |
|  |  |  | Raw material – GWLM | 1mL/lot | <1CFU/mL |
|  |  |  | Raw material - FSM | 1mL/lot | <1CFU/mL |
|  | Total Yeast and Mold count (TYMC) |  | Final container Working Cell Bank | 1mL | <1 CFU/mL |
|  |  |  | Raw material – GWLM | 1mL/lot | <1CFU/mL |
|  |  |  | Raw material - FSM | 1mL/lot | <1CFU/mL |
|  | Microbial contamination test |  | Final container Working Cell Bank | 1mL/microbe tested except Salmonella.  Salmonella: 10mL  (Total: 14mL) | Absence of bile-tolerant Gram negatives, *Staphylococcus aureus*, *Pseudomonas aeruginosa*, *Escherichia coli*, and Salmonella |

WCB: Working Cell Bank. AST: Antimicrobial Susceptibility Test. gDNA: Genomic DNA GWLM: Graver-Wade liquid medium. FSM: Freezer storage medium. CFU: Colony forming unit.

**Table S4. Dose Cell Bank product specification sheet**

| **Test category** | **Test** | **Accreditation** | **Sample tested** | **Amount of sample tested (minimum)** | **Specifications** |
| --- | --- | --- | --- | --- | --- |
| Release tests: Dose determination | CFU enumeration | Not accredited | Final dose suspension with cryopreservative (prior to freezing) | Each dose level enumerated in technical triplicate | Report only |
|  |  |  | Final container DCB vials (thawed) | 3 vials per dose level | 10^8^ ± 0.5log_10_  10^7^ ± 0.5log_10_  10^6^ ± 0.5log_10_  10^5^ ± 0.5log_10_  10^4^ ± 0.5log_10_ |
| Release tests: Culture identification | MALDI | ISO 15189 | Final container dose cell bank | Per dose level: 3 colonies/ 1 DCB vial | *N. gonorrhoeae* |
|  | Gram stain |  | Final container dose cell bank | Per dose level: 3 colonies/ 1 DCB vial | Gram negative diplococci with no visible contamination |
|  | Superoxol test |  | Final container dose cell bank | Per dose level: 3 colonies/ 1 DCB vial | Positive |
|  | Oxidase test |  | Final container dose cell bank | Per dose level: 3 colonies/ 1 DCB vial | Positive |
|  | Nucleic acid amplification test (NAAT) | Not accredited for this matrix | Final container dose cell bank | 1 vial (10^4^ dose) in triplicate. 1 vial 10^8^ dose) in triplicate. | Positive |
| Release tests: AMR characterisation | Antimicrobial susceptibility:  Penicillin  Spectinomycin  Ceftriaxone  Ciprofloxacin  Tetracycline  Azithromycin | ISO 15189 | Final container dose cell bank | 1 vial, 3 colonies (10^8^ CFU/mL dose vial) | Penicillin: susceptible or less susceptible  Spectinomycin: susceptible  Ceftriaxone: susceptible  Ciprofloxacin: susceptible  Tetracycline: susceptible  Azithromycin: likely susceptible |
| Release tests: Genomic identification and characterisation | Whole genome sequencing | Not accredited | Working Cell Bank gDNA | gDNA extract from DCB vial | MLST: 1596  NG-STAR: 4332  NG-MAST: (*tbpB* 3072, *porB* 7697) |
| Release tests: Microbiological limits and material sterility | Total aerobic microbial count (TAMC) | Accredited for Appendix XVI B, British Pharmacopeia. | Final container dose cell bank | 1mL (2 vials) per dose level | Report only |
|  |  |  | Raw material (GWLM) | 1mL/lot | <1CFU/mL |
|  |  |  | Raw material (FSM) | 1mL/lot | <1CFU/mL |
|  | Total Yeast and Mold count (TYMC) |  | Final container dose cell bank | 1mL (2 vials) per dose level | <1 CFU/mL |
|  |  |  | Raw material (GWLM) | 1mL/lot | <1CFU/mL |
|  |  |  | Raw material (FSM) | 1mL/lot | <1CFU/mL |
|  | Microbial contamination test |  | Final container dose cell bank | 2 vials (1mL) per microbe tested (except Salmonella) per dose level  Salmonella: 20 vials (10mL) per dose level  Total: 28 vials (14mL) per dose level | Absence of bile-tolerant Gram negatives, *Staphylococcus aureus*, *Pseudomonas aeruginosa*, *Escherichia coli*, and Salmonella |
| Characterisation: colony morphology | Opa characterisation | Not accredited | Final container dose cell bank | Minimum 1 DCB vial | Report only |
|  | Pili characterisation | Not accredited | Final container dose cell bank | Minimum 1 DCB vial | Report only |

DCB: Dose Cell Bank. AST: Antimicrobial Susceptibility Test. gDNA: Genomic DNA GWLM: Graver-Wade liquid medium. FSM: Freezer storage medium. CFU: Colony forming unit. GWLM: Graver-Wade liquid medium. FSM: Freezer storage medium.

**SUPPLEMENTARY FIGURES**

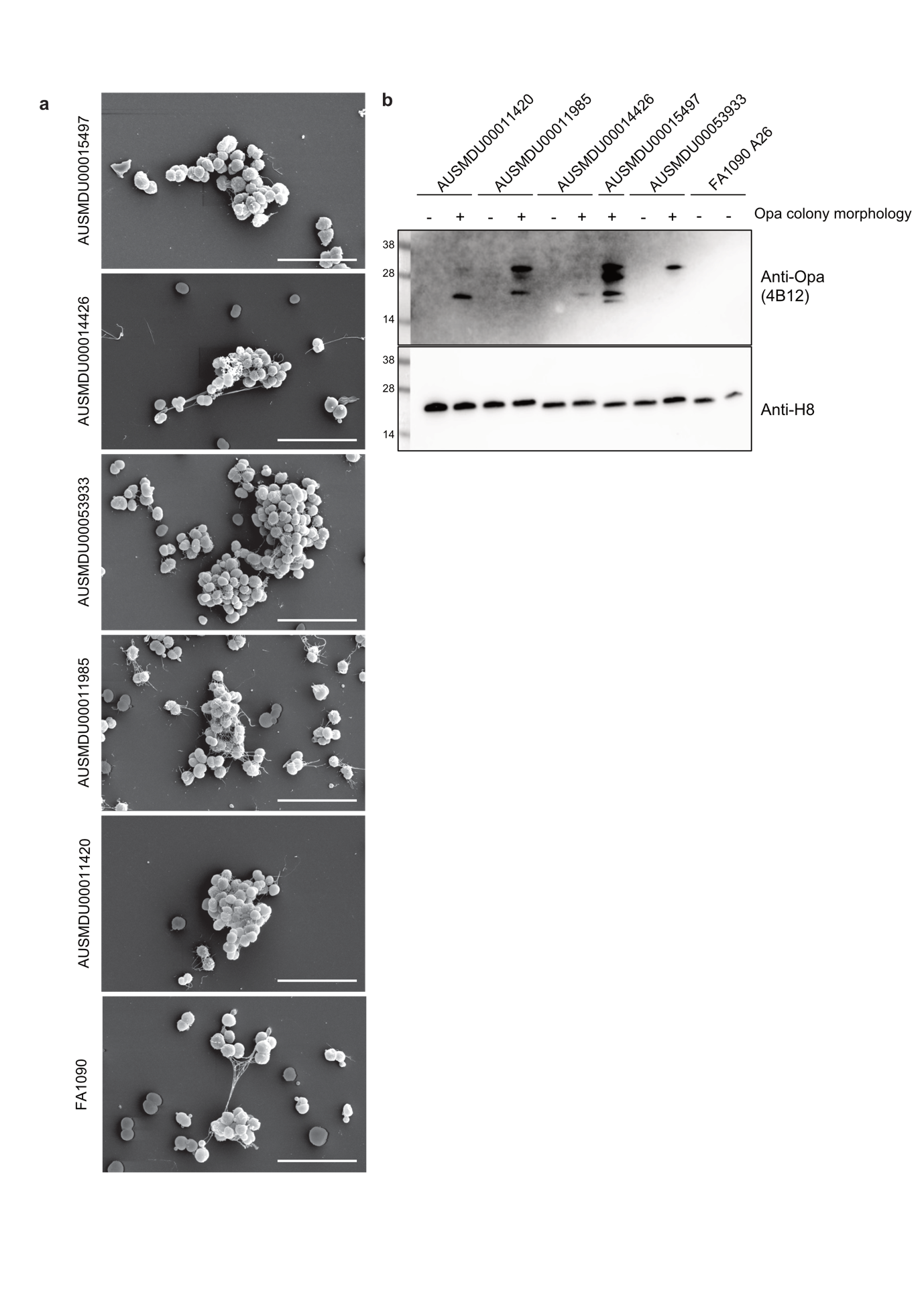
**Figure S1. Candidate *N. gonorrhoeae* isolate morphology**

(a) Representative scanning electron micrographs of *N. gonorrhoeae* isolates mounted onto glass coverslips, imaged at 25,000x magnification, scale bar represents 5μm. (b) Representative immunoblot showing Opa expression in Opa- and Opa+ colony variants of each isolate, H8; loading control.

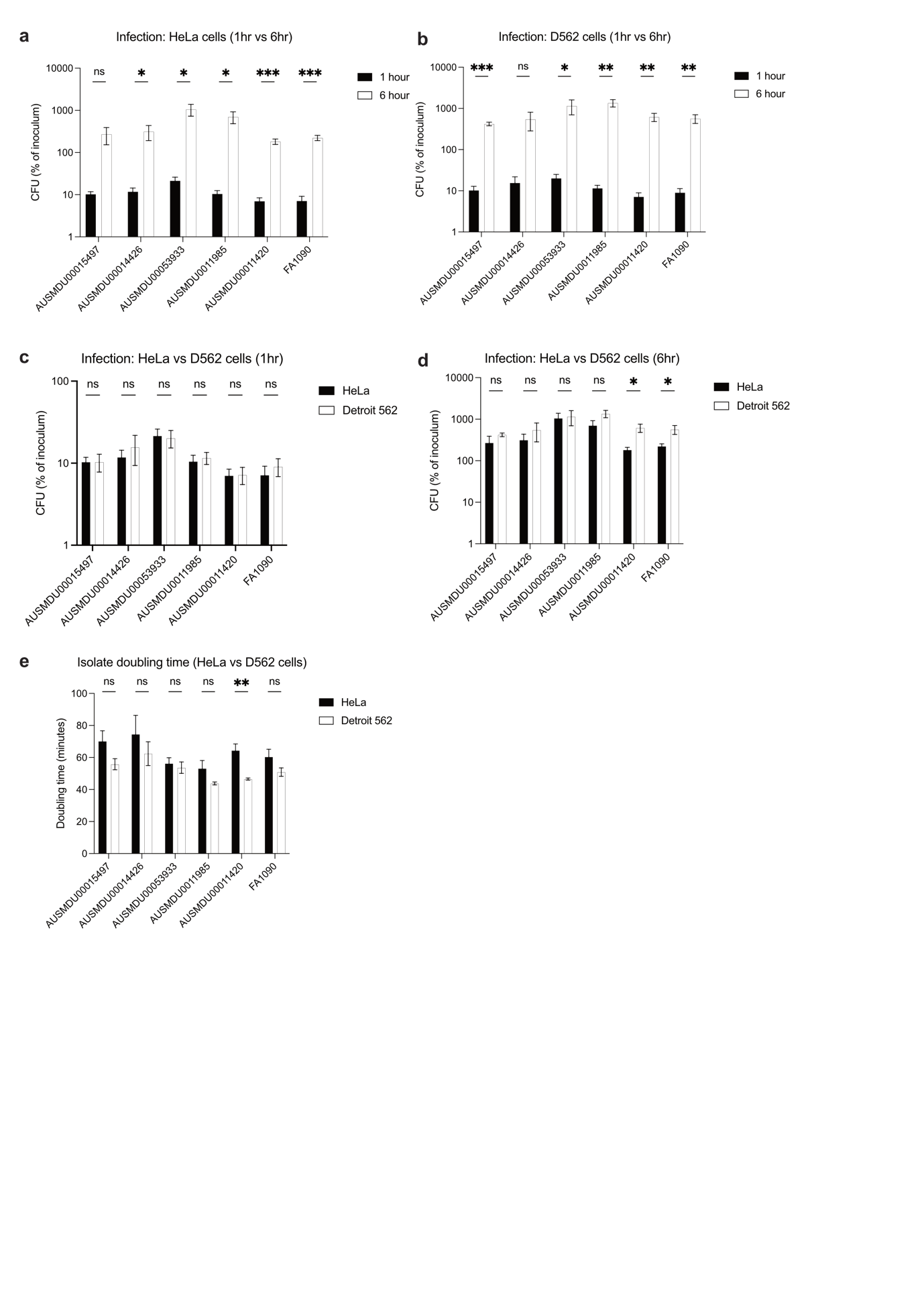
**Figure S2. Comparison of infection dynamics in HeLa and Detroit 562 cells**

(a-b) Comparison of CFU at 1 hour and 6 hours post-infection of (a) HeLa cells or (b) Detroit 562 cells for *N. gonorrhoeae* candidate isolates. Results are presented as a percentage of the input (inoculum). (c-d) Quantification of CFU from infected HeLa cell and Detroit 562 cells at (c) 1hr post-infection and (d) 6 hours post-infection. Results are presented as a percentage of the input (inoculum). (e) Doubling time in minutes of each *N. gonorrhoeae* isolate during 6-hour infection of HeLa and Detroit 562 cells. Bar and error represent mean ±SEM. Significance was calculated using *t-*test *P<0.05, **P<0.01, ***P<0.001, ns not significant.

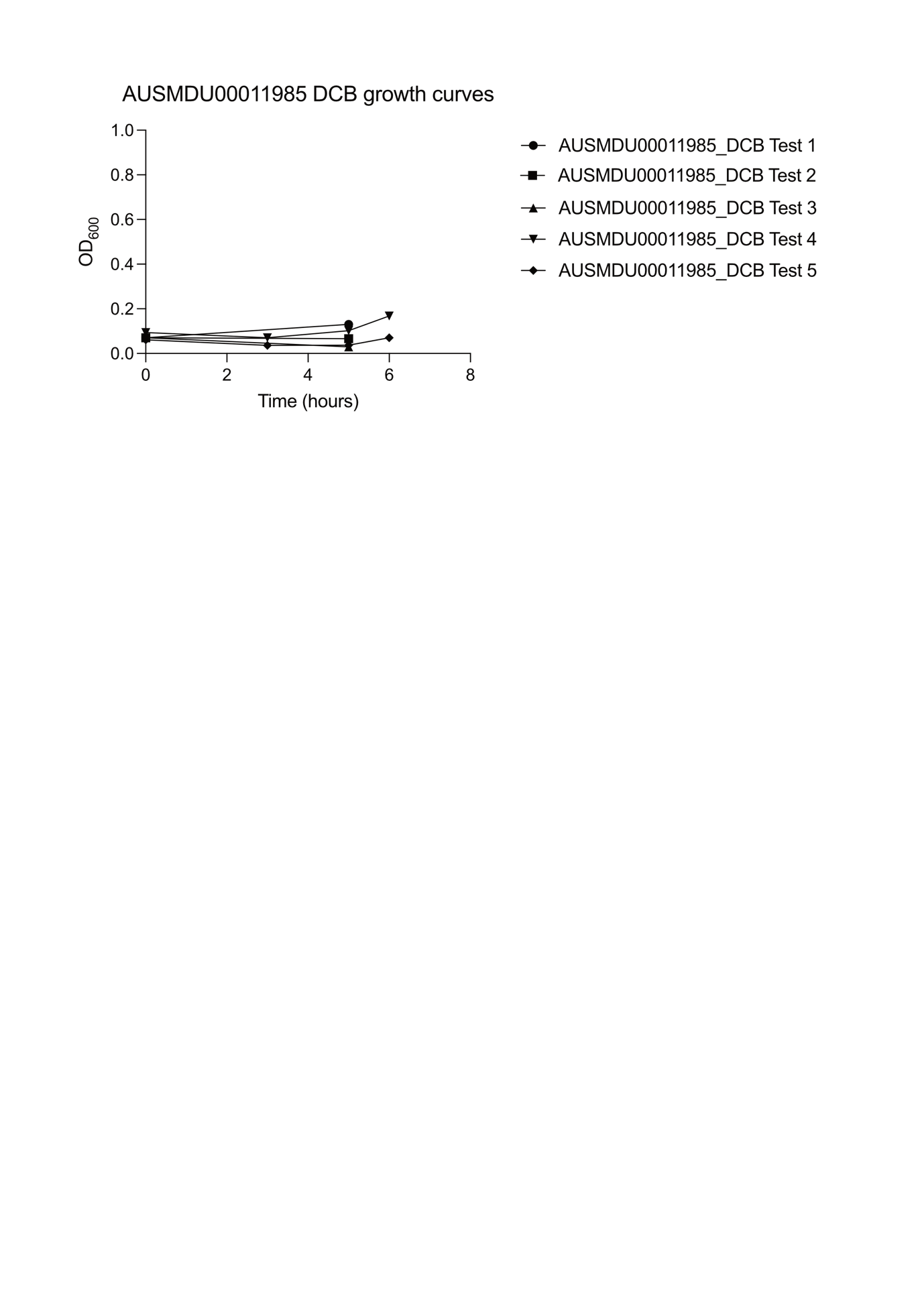
**Figure S3. Assessment of AUSMDU00011985 manufacture performance**

Growth curves (OD_600_) of AUSMDU00011985 in the liquid growth phase of Dose Cell Bank (DCB) manufacture.

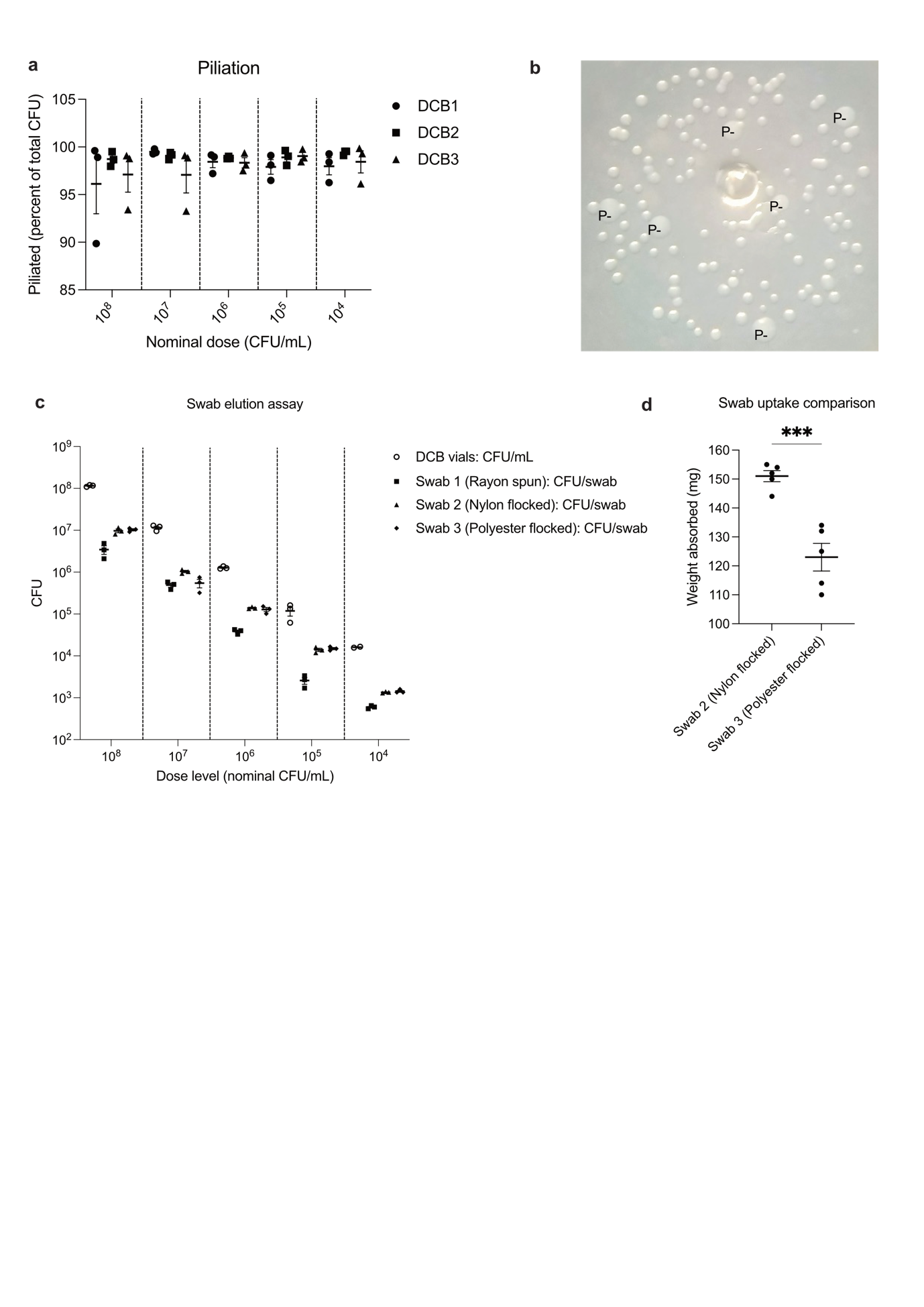

**Figure S4. *N. gonorrhoeae* AUSMDU00053933 Dose Cell Bank characterisation and delivery**

(a) Percentage of *N. gonorrhoeae* AUSMDU00053933 colonies from DCB vials deemed piliated based on large, flattened appearance. Filled circles represent average piliation percentage from each Dose Cell Bank family, line and error represent mean ±SEM. (b) Representative image of *N. gonorrhoeae* AUSMDU00053933 colonies demonstrating identification of non-piliated colonies (P-) based on large, flattened appearance as viewed using dissecting microscope.

(c) Enumeration of *N. gonorrhoeae* AUSMDU00053933 CFU directly from DCB vials (CFU/mL) or eluted from three different swab types (CFU/swab) after submersion in DCB vials. Filled circles represent individual datapoints, line and error represent mean ±SEM. (d) Comparison of swab liquid uptake, measured in milligrams, for two swab types. Filled circles represent individual datapoints, line and error represent mean ±SEM. Significance was calculated using *t-*test, ***P<0.001.
